# Cold trapped - Correcting locomotion dependent observation biases in thermal preference of *Drosophila*

**DOI:** 10.1101/400069

**Authors:** Diego Giraldo, Andrea K. Adden, Ilyas Kuhlemann, Heribert Gras, Bart R. H. Geurten

**Author notes:** authors contributed equally (Diego Giraldo, Andrea K. Adden).

## Abstract

Sensing environmental temperatures is essential for the survival of ectothermic organisms. In *Drosophila,* two methodologies are used to study temperature preferences (T_P_) and the genes involved in thermosensation: two-choice assays and temperature gradients. Whereas two-choice assays reveal a relative T_P_, temperature gradients can identify the absolute T_p_. One drawback of gradients is that small ectothermic animals are susceptible to cold-trapping: a physiological inability to move at the cold area of the gradient. Often cold-trapping cannot be avoided, biasing the resulting T_P_ to lower temperatures. Two mathematical models were previously developed to correct for cold-trapping. These models, however, focus on group behaviour which can lead to overestimation of cold-trapping due to group aggregation. Here we present a mathematical model that estimates the behaviour of individual *Drosophila*in temperature gradients. The model takes the spatial dimension and temperature difference of the gradient into account, as well as the rearing temperature of the flies. Furthermore, it allows quantifying cold-trapping, reveals true T_P_, and differentiates between temperature preference and tolerance. Online simulation is hosted at http://igloo.uni-goettingen.de. The code can be accessed at https://github.com/zerotonin/igloo.

## Introduction

In recent years, the thermoreceptive system of *Drosophila* has been the subject of intense study: Starting with the discovery of the role of the *painless* gene by ^1^, a number of receptor genes (*painless*^2,3^; *pyrexia*: ^4^; *dtrpA1*:^5,6 5,6^; *dtrp, dtrpL*: ^7^; *brv*:^8^; Gr28b.d: ^9^), signalling pathways (cAMP-PKA-pathway:^10^; phospholipase C pathway:^11^) and brain regions ^12–14^have been shown to be involved in thermosensation and the processing of temperature information.

Most of the studies used behavioural assays to assess gene and protein function in thermosensation. These behavioural tests are used to determine the temperature preference (T_P_) and generally belong to one of two types: either a two-choice assay (^15^; e.g. ^7,8^) or a temperature gradient (^15^; e.g. ^4,5,16^.). In a two-choice assay, flies are given the choice between a “preferred” and a “non-preferred” temperature. Such assay is useful to establish a relative T_P_ and to measure avoidance behaviour, but the paradigm is less suited for establishing absolute T_P_. An absolute T_P_ can be determined in a temperature gradient: animals are introduced into a gradient ranging from cold to hot temperatures, and are allowed to move within the gradient in order to reside at the T_P_. One of the drawbacks of this paradigm is that animals may be cold-trapped in the low temperature range. Since *Drosophila* is an ectothermic organism, its locomotion speed critically depends on the ambient temperature (T_A_). Thus, when the flies traverse colder regions, they slow down and linger there, and can fall into a stasis-like state ^17^. This velocity reduction will bias the identification of T_P,_ as an observer will underestimate T_P_, because flies will reside for longer period in colder temperatures due to their reduced ability to leave those regions. To avoid this bias one can correct the resulting data by comparing it to a null model. Such a null model would represent an animal without T_P_ whose locomotion depends on the ambient temperature.

Null models for temperature-dependent locomotion have been developed for cohorts of small ectotherms (*Caenorhabditis elegans*: ^18^, *Drosophila melanogaster*: ^19^). Cohorts add an ethological bias by group aggregation. Aversive stimuli are avoided more strongly in groups than in single individual ^20^ and sensing as well as decision making are enhanced in groups ^21,22^. Both factors will bias the individual sensation and eventually the T_P_ of a fly. Especially in spatial tasks (e.g. positioning within a temperature gradient) individual-based-models (IBM) surpass group models ^23^.

Using a large behavioural database of approx. 10 million walking-velocity combinations of *D. melanogaster* larvae and adults measured at different temperatures, we formulated a null model named IGLOO (IGLOO is a <u>G</u>radient <u>LO</u>comotion m<u>O</u>del) that extends existing models. The model we propose allows predicting individual fly trajectories in dependence of the ambient (T_A_) and rearing temperature (T_R_). We show that T_R_ strongly influences locomotion at different T_A_, and offer a null model to correct for these and other biases introduced e.g. by cold-trapping. IGLOO distinguishes between absolute T_P_, tolerance and avoidance of temperature, reducing the number of experimental trials and tested animals. IGLOOs most striking feature is its ability to predict if a mutant phenotype arises on the level of internal or external thermosensory systems.

## Results

Our model is a simple random walk model (review ^24^). In general, Random walk models include an agent that moves in steps of random direction through a virtual environment. Our environment is a one-dimensional temperature gradient in which the temperature between both extremes is linearly distributed. Therefore, our agent, the simulated fly, can only move up or down the gradient. Each step of our agent represents a bout of walking for the fly. The step size is derived from the step velocity and the step duration, which both depend on body temperature (T_B_). The fly’s TB is calculated by the ambient temperature (T_A_ | see Eq. 4). In pseudo code, our model works as follows (compare Fig. 1):

**Figure 1:**
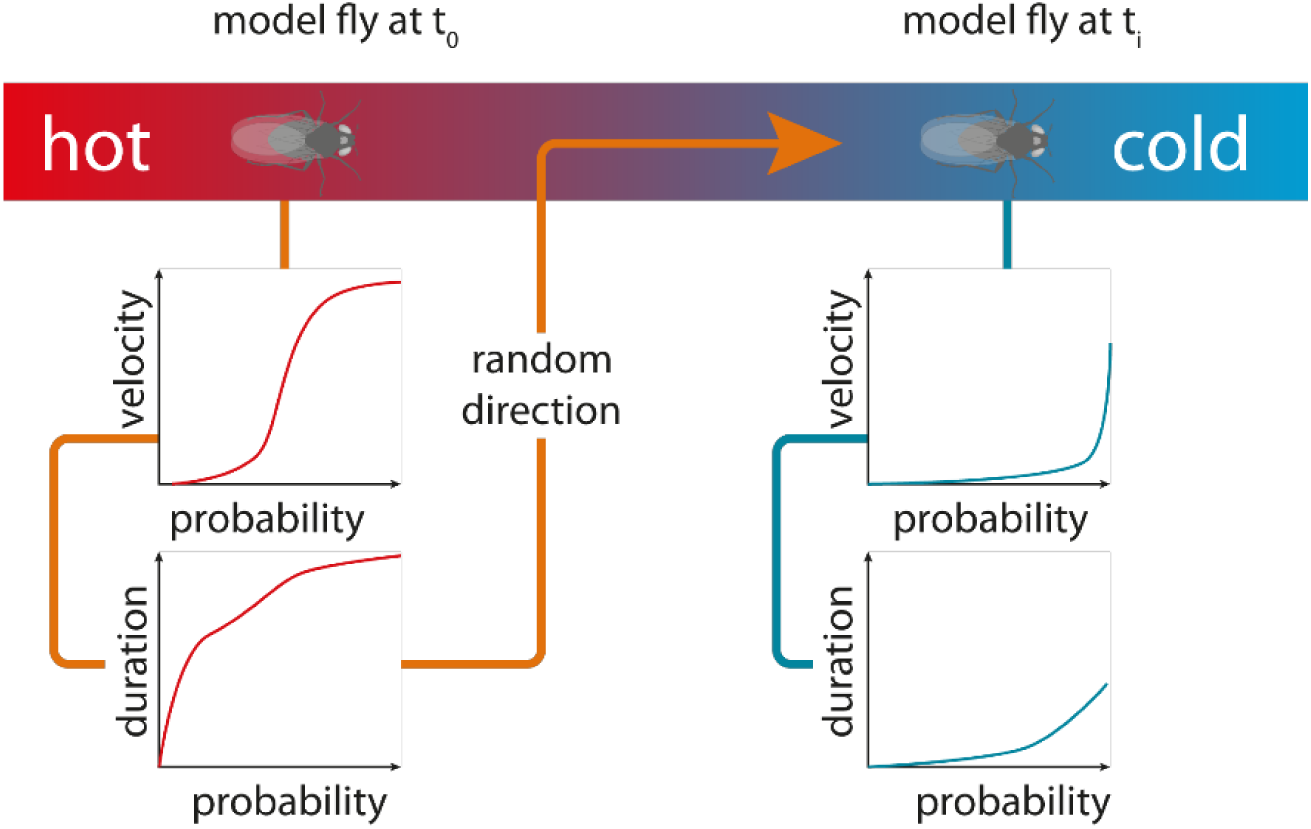
Model Schematic. The red to blue bar represents the temperature gradient, with extremes as indicated. The simulated fly at time t_0_ is positioned at the warmer end and therefore its probability functions would more often render fast velocities and long bout durations than at time point t_i_. Graphs do not show real data and omit the influence of the rearing temperature (T_R_).

1. Update body temperature from ambient temperature (Eq 4., see Methods)
2. Pick a random number (0.0 to 1.0) = P_V_
3. Calculate a step velocity from histogram fit using P_V_ (Eq 2.2, see Methods)
4. Pick a random number (0.0 to 1.0) = P_D_
5. Calculate step duration from histogram fit using P_D_ (Eq 3.2, see Methods)
6. Pick random direction (50% + to 50% -) = P_Dir_
7. Make Step

Note that this can create steps that could make the agent (fly) collide with the gradient extremes. In these cases, the walls reflect the animal, and the animal instantly reverses its direction. This is equivalent to an infinite environment with repetitive triangular temperature gradients. Phases 1 to 7 are repeated until the accumulated step duration exceeds the simulation duration. Then the trace of the agent through the environment can be calculated as well as probability density in the temperature gradient.

The model is based on Benzer climbing assays of flies reared at different temperatures (T_R_) and larvae moving at different T_A_. Histograms of the velocity and bout duration of over 400 individual flies in total allow building a probability function for a bout of walking at a given temperature. Each individual bout has a certain velocity and duration, based on the T_B_ and the T_R_ of the modelled individual. We accumulated over 24 million individual velocity frames and found qualitative and quantitative differences in the locomotion behaviour depending on the T_A_ (Fig. 2). A clear influence of the T_A_ on the median speed was observed (Fig 2A). Larvae (wt_L_) and adults showed almost no movement at very low ambient temperatures, and their median speed increased with T_A_. Wild-type adults raised at 30°C (wt_30_) were generally slower (Fig. 2A) and, compared to adults, larvae showed a higher average velocity. It should be noted that wt_30_ flies were generally less active as well as viable and that a large proportion of their larval population died before or during pupation.

**Figure 2:**
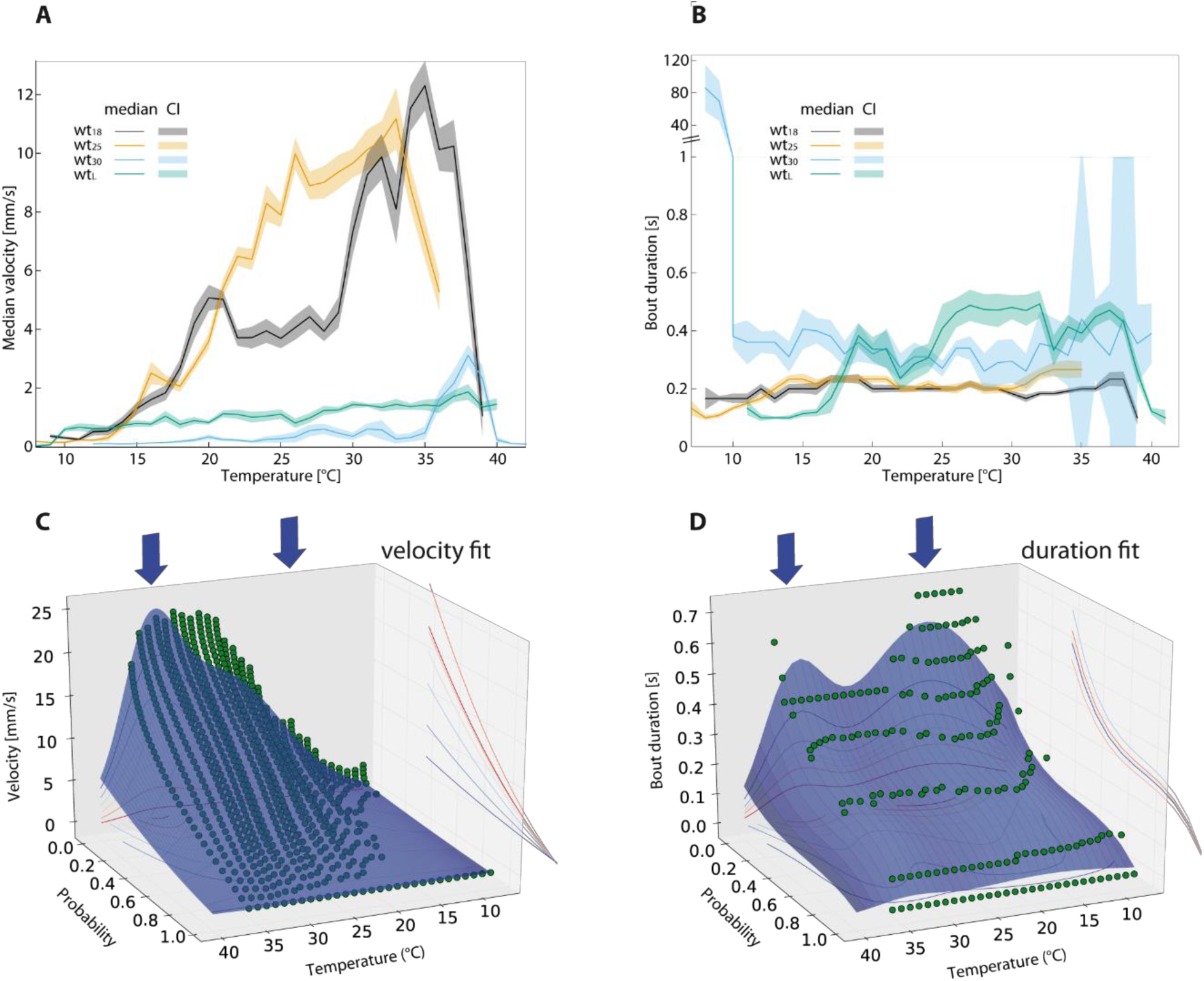
Activity and walking speed of adult flies at different temperatures. A) Median velocity in mm/s plotted against the ambient temperature. The solid line demarks the median. The shaded area around it marks the 95 % confidence interval (n = 50 animals per sample). B) Locomotion bout duration plotted against temperature as in A. C) Probability for a velocity to occur at different ambient temperatures in wt_25_ (green dots). The shaded area below is the result of the velocity fit function (see Method section), lines on the wall are the respective projections of the shaded area onto this axis. The blue arrows above the plots denote values of the two Gaussian functions used to fit the temperature domain of the fit-function: = 26°C and = 34°C. D) Bout duration fit function shown as in C _2_(_5_w).t Additional fit functions can be found in the Supp. Mat.

The second attribute our model is based on is the walking bout duration. This duration is constant over a large T_A_ range for wild-type adults raised at 18°C (wt_18_) or at 25°C (wt_25_), only wt_L_ reduce their bout duration significantly below 18°C T_A_ and above 38°C T_A_ (Fig. 2B).

wt_18_ were more active at lower T_A_ than wt_25_ or wt_30_, whereas the opposite effect was seen at higher T_A_ (supp. Fig. 1). The activity index is the fraction of time the animal spent moving. wt_18_ and wt_25_ are able to sustain high activity levels throughout a large part of the T_A_ spectrum, while wt_30_ has a more erratic activity pattern (see discussion). wt_L_ show very little activity in the lowest part of the range but if T_A_ was above 10°C the activity index (see materials and methods) increase up to 60%. wt_L_ seems unaffected by the hottest ambient temperatures of the spectrum, unlike adults (supp. Fig. 2D).

To make the model simulation faster we fitted functions to the histograms as shown for wt_25_ in Fig. 2 C,D and for other strains in supp. Fig. 2. All further calculations shown in this article were also undertaken on the respective original histogram, qualitatively and quantitatively producing nearly identical results.

One of the most striking direct results was the emergence of similar (=central value of the Gaussian distribution | σ width of the Gaussian distribution) in the temperature Gaussian functions of the fits to velocity and duration. As can be seen in Fig. 2C,D (arrows), our model centres these Gaussian functions for wt_25_ at μ = 26 °C and σ ≈ 5.41 °C, and μ = 34 °C and σ = 2.27, respectively (temperature and bout duration). These temperatures resemble the thermal optima for transduction channels in thermosensation (dTRPA1: 25°C ^5^) and nociception (>38°C ^2,4,25^) of adult flies.

To simulate the probability density of a fly without T_P_ in a thermal gradient we used the aforementioned equations in a random walk paradigm. In the initial state the fly’s T_B_ equals the T_A_. The model uses the interpolated probability function to determine the velocity and duration of a bout based on T_B_. The direction of the bout is random. T_B_ is updated with respect to T_A_ via the thermal transduction equation described in the Methods section, functionally equalling a low pass filter (Fig. 3A).

**Figure 3:**
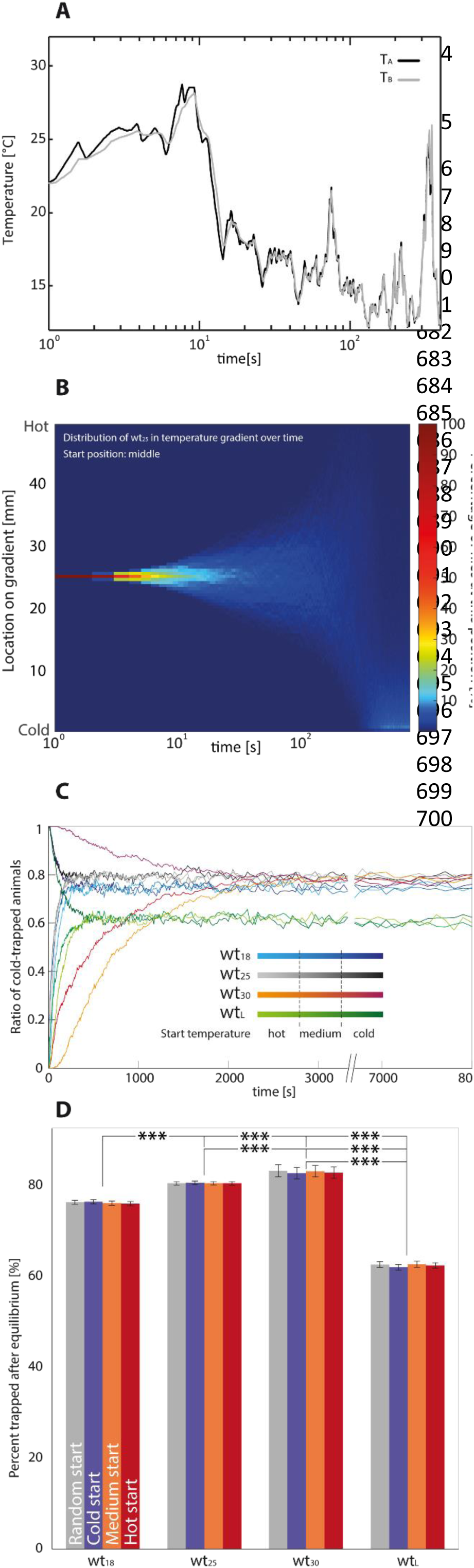
Model evaluation. A) The body temperature of a single simulated fly (adult raised at 25°C) is plotted over time (grey line). The body temperature (T_B_) differs from the ambient temperature (T_A_) (black solid line) according to the heat conduction equation (eq. 4). B) Position of all 1000 simulated flies (wt_25_) as a histogram over time, with the position on the y-axis, time on the logarithmic x-axis and percentage of animals colour-coded as shown on the right. All animals started in the middle of the temperature gradient and then began their random walk. At the lower right corner one can see a small but bright line, which is indicative of a stable equilibrium of cold-trapped animals. C) Ratio of cold-trapped simulated animals plotted over time. An animal is considered cold-trapped when it is in the coldest 4 mm of the gradient. Colours code for different rearing situations, the shade of the colour denotes the temperature the simulation was started on the gradient. D) Median percentage of cold-trapped individuals after equilibrium stabilised. Values are median ± confidence interval of the median. Colour code indicates starting position. *** = p < 0.001; Fisher’s permutation test, corrected with Benjamini-Hochberg FDR procedure.

We simulated one thousand flies to obtain a null-probability density of a fly without T_P_ (e.g. Fig. 3B). As can be seen in figure 3C, after about 10 minutes (50 min for wt_30_) a stable ratio is formed in which a large percentage of the flies aggregate at the cold end of the gradient, apparently being cold-trapped. In a modelled temperature gradient ranging from 14°C to 32°C, the probability of cold-trapping, as well as the overall distribution within the gradient (see Fig3B, supp. Fig.3 and sup. Fig4), depend on T_R_ of the flies and the developmental stage (Fig. 3D). When starting from a medium or high temperature, however, all strains take equally long to become cold-trapped for the first time (Fig. 3C), with the exception of wt_30_ and wt_L_, which took much longer due to their slow speed of locomotion (compare Fig. 2A and 3C). We found that after running the simulation for 10 minutes, a stable distribution of approximately 70-80% cold-trapped adult flies and approximately 60% cold-trapped larvae was always established (Fig. 3D), except of wt_30_ which needed 60 min. wt_18_ flies were less frequently cold-trapped than wt_25_ and wt_30_. However, the distributions of non-cold-trapped flies were non-linear in the warmer range of the gradient and differed between T_R_ cohorts.

By subtracting the modelled null distribution from the measured distribution of adult flies in a temperature gradient, avoidance, T_P_ and tolerances can be easily identified and visualized (see Fig. 4 A-C). As a result of using the 95% confidence interval of the median (CI) there is no significant preference or avoidance of the respective temperature when the CI of measured data includes the modelled data, otherwise there is (p< 0.05). Furthermore, there are values that are not significantly avoided or preferred at the fringes of the tolerable temperature zone (grey value Fig. 4 A-C). Our analysis allows for a more differentiated interpretation of temperature behaviour.

**Figure 4:**
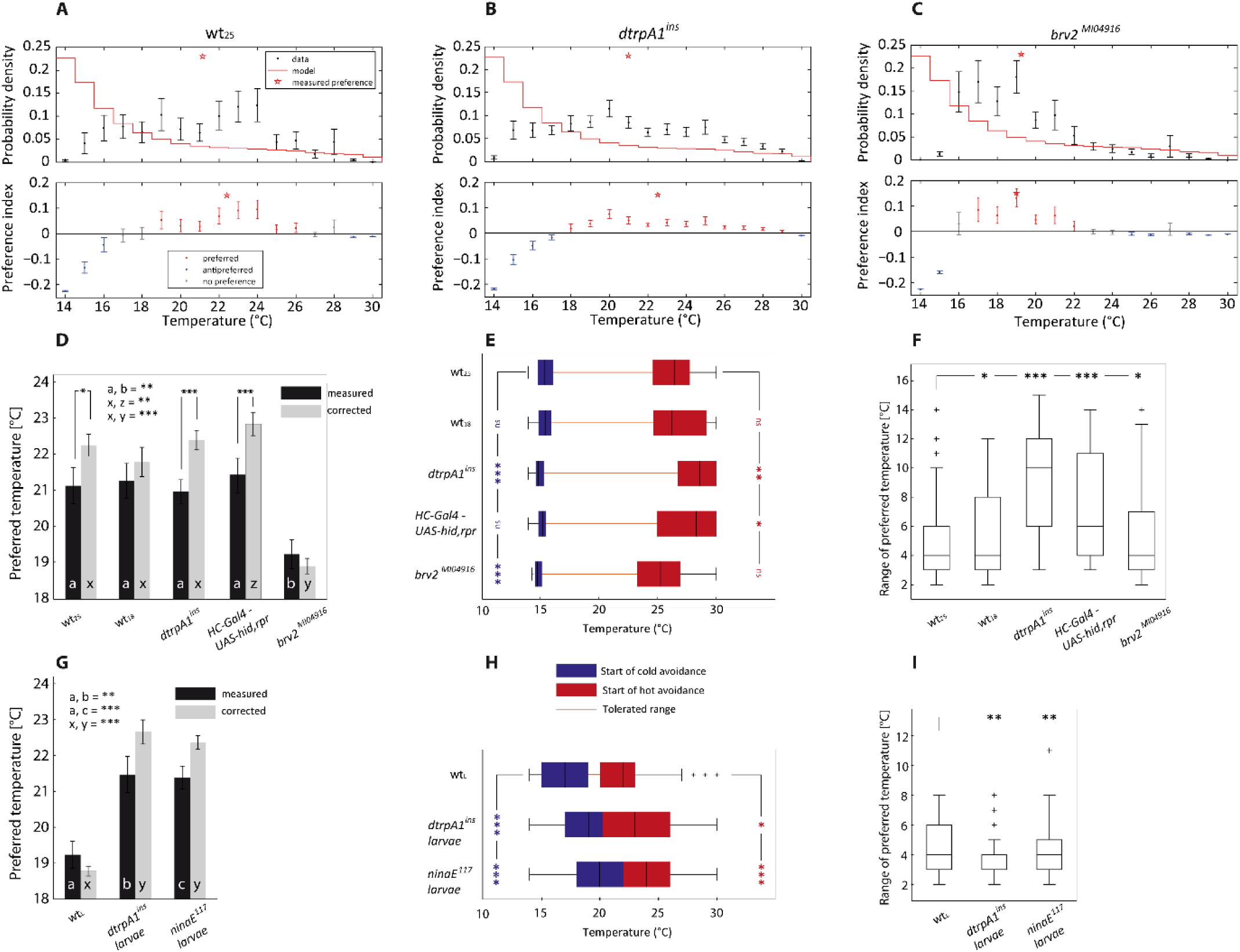
Correction of results gathered from different Drosophila strains in experiments with a temperature gradient. A-C) Upper row shows raw density distribution of animals in a 14°-30°C temperature gradient. Values are median ± 95% confidence interval of the median. The red star indicates the T_P_ of the tested animals. The red stair plot depicts the result of simulated flies. The lower plot shows the resulting values after corrections (Preference index = prob_observation_ - prob_null model_). Red: temperatures that are significantly more preferred than the null model would predict. Blue: temperatures that are significantly less preferred. Grey: data that is not significantly different from the null model. The error bars represent the 95% confidence interval of the median. (A: wt_25_ n = 45 | B: *dtrpA1*^*ins*^ *n= 47* | C: *brv2*^*MI04916*^ *n= 37)* D) Preferred temperatures before and after correction by IGLOO. Plotted are medians ± 95% confidence interval. Significances are shown either by stars or letters (indicated in graph). Comparisons before and after correction where done by a paired t-test and corrected with Benjamini Hochberg FDR. Comparisons between groups were done with a normal t-test and again corrected via Benjamini Hochberg FDR (n-values for D-F: wt_25_ n = 45 | wt_18_ n = 50 | *dtrpA1*^*ins*^ *n= 47* | *HC-Gal4 – UAS-hid,rpr* = 40 | *brv2*^*MI04916*^ *n= 37*). E) Cold avoidance as blue boxplot and hot avoidance as a red boxplot. The median is indicated through black vertical line separating the upper and lower quartile. Whiskers mark the 1.5 interquartile distance. The line connecting both boxplots (orange) indicates the tolerable temperatures. Stars on the left side in blue indicate significant differences to wt_25_ of the cold avoidance median. Significances were detected using Fisher’s Permutation test, corrected as before. The red stars on the right side indicate significant differences to wt_25_ of the median hot avoidance, in identical fashion. F) Range of the preferred temperatures plotted as boxplots, with definitions as in E. Note that the preference range shows the expanse of temperature which are categorised as preferred in each individual fly. Hence even though red and blue boxes of E (wt_25_) are more >10°C apart the median preference range is only 4 °C wide, as the difference could be values that are insignificantly different from the null model. Stars again show significantly different values in comparison to wt_25_ (Fisher’s permutation test, Benjamini Hochberg FDR). G-I) show equivalent plots to D-E for larvae data (n-values for G-I: wt_L_ =72 | *dtrpA1*^*ins*^ = 60 | *ninaE*^117^ *=*74).

To test the model, we assessed the T_P_ of wt_18_ and wt_25_ in a temperature gradient and corrected the measured distribution. After the correction, we calculated the new T_P_ (Fig. 4D), the start of cold and hot avoidance, and the range of tolerated temperatures. The mean T_P_ was increased after the correction in both groups tested, proving that cold-trapping biases T_P_ and that a correction of T_P_ is needed. The change in T_P_ was significant for wt_25_ but not for wt_18_, suggesting that wt_25_ are more susceptible to cold-trapping. The median T_P_ obtained for wild-type (22.5°C) corresponds well to already published T_P_ ^15,26^ when taking the low humidity in the temperature gradient into account ^27^. Although wt_18_ distributed more broadly in the gradient than wt_25_ (supp. Fig. 5), the overall T_P_, and the start of hot and cold avoidance were similar for both groups. This suggests that although there is an effect of T_R_ on locomotion speeds (Fig. 2A), this in fact does not affect T_P_ (Fig. 4D).

As a proof of concept we tested two fly strains with impaired heat sensation, i.e. flies with ablated hot sensing neurons in the arista (hot cells) and a genome-wide knock down of the heat sensitive channel dTRPA1 (*dtrpA1*^*in*^^*s*^ mutants). We also chose a strain with impaired cold sensation (i.e., *brv2*^*MI04916*^ mutant flies lacking Brivido2, a channel involved in cold sensation ^8^) to test if IGLOO would reveal their T_P_ faithfully, without masking the original phenotype. To ablate hot cells (HC) we expressed the apoptotic factors Hid and Reaper ^28,29^ in HC using a *HC-Gal4* driver line. Both HC-ablated and *dtrpA1*^*ins*^ have a well-characterised heat-sensing impairments ^5,8^.

The median T_P_ was significantly increased in *drtpA1*^*ins*^ after correction, however the resulting median T_P_ was not significantly increased compared to wt_25_ controls (Fig. 4D). Nonetheless both the start of hot avoidance (Fig. 4E) and the range of tolerated temperatures (Fig. 4F) were significantly increased in these flies. The start of cold avoidance was significantly reduced, indicating that these flies have also problems in cold temperature avoidance.

The median T_P_ of HC ablated flies was also significantly increased after correction. In contrast to *drtpA1*^*ins*^ flies, the resulting median T_P_ was significantly higher than in wt_25_ controls (Fig. 3D). This difference was not observed before correction, indicating that cold-trapping was obscuring the hot sensing impairment. The start of hot avoidance was significantly increased, whereas the cold avoidance was unaffected (Fig. 4E). The T_P_ of these flies had a wider range of T_P_ distribution than wt_25_ controls which was shifted toward warmer T_A_ (Fig. 4F), indicating that the flies were impaired in heat sensation as expected.

The mean T_P_ of *brv2* ^*MI04916*^ mutants was not significantly different after correction compared to the uncorrected T_P_, but was significantly lower than in wt_25_ controls (Fig. 4D). Additionally, the start of cold avoidance was significantly reduced, whereas the start of hot avoidance was unaffected (Fig. 3E). The overall distribution of these flies was shifted towards colder temperatures, suggesting an impairment in cold sensation. The fact that a cold impairment is still observed after correction shows that correction with IGLOO still allows for the detection of mutants defective in cold sensing.

wt_L_ had a median T_P_ around 18.5°C and did not show a significant change in the median T_P_ after correction, suggesting that cold-trapping is less severe for larvae than for adults (Fig. 4G). This is consistent with the temperature dependence of locomotion (Fig. 2A,B). The start of hot avoidance was around 25°C, and the start of cold avoidance around 16°C (Fig. 4H). In contrast to their wild-type conspecifics, *dtrpA1*^*ins*^ larvae had a significant increase in the T_P_ after correction, and their T_P_ was significantly higher than that of wt_L_. Additionally the start of cold and hot avoidance was significantly increased (Fig. 4 H,I). The overall distribution of the mutant larvae in the gradient was shifted to warmer temperatures, suggesting that these larvae are impaired in heat sensation. A similar effect as in *dtrpA1* ^*ins*^ larvae was observed in *ninaE*^*17*^ mutants lacking Rhodopsin1, which is supposed to interact with TRPA1 channels and to be indispensable for warm temperature sensation^27^.

## Discussion

In this article we provide an individual-based-model for temperature gradient locomotion in small ectothermic animals. Especially in ectotherms such as *Drosophila*, the ability to move strongly depends on T_A_ – a fact that may bias experimental results. Two pioneering models that address the problem were published by Anderson et al. (2007) and Dillon et al. (2012). Both are diffusion-based models that simulate cohorts of animals in a thermal gradient and produce overall animal distributions. This is useful for experiments comparing e.g. the resulting null-distribution to a distribution obtained in an experiment. The drawback of group behaviour is the bias caused by group aggregation ^20–22^, which influences sensation, decision making and enhances synchronisation of behaviour. Therefore, it is not surprising that for locomotion Individual-Based-Models are preferred ^23^. Given that many studies on thermosensation use group behaviour instead of individual behaviour ^5,8,9,15,19^, but differences between group and individual studies in gradient behaviour are regularly observed, the potential choice of individual T_P_ over group T_P_ could become an important consideration when designing an experiment. In any case, modelling single-fly trajectories is essential for a detailed analysis of the behaviour of individual flies. We therefore present a simple model that is based on data of walking *Drosophila* at various temperatures.

T_A_ strongly influences the locomotor performance in both larvae and adults. The strength of this influence, however, was much smaller in larvae, which revealed an almost linear relation between temperature and speed. These results suggest that larvae may have physiological mechanisms that make them more resistant to cold, probably because they cannot escape as fast as adults. To confirm this, when the experiments were carried out in the gradient to assess T_P_, larvae never seemed to enter a chill coma like the adults. Nevertheless, a cold bias may compromise the uncorrected larval gradient data since cold temperatures will make larvae slower, making the correction still necessary.

Rearing temperature also influenced the resistance to cold temperatures. Adults reared at 18°C (wt_18_) were more resistant and had higher activity in colder temperatures than adults reared at 25°C (wt_25_) which also resulted in a lower proportion of cold-trapped animals in the wt_18_ simulation. The resistance to cold temperatures in wt_18_ is consistent with previous studies demonstrating that development at low temperatures reduces the chill coma temperature and increases the resistance to cold in different *Drosophila* species ^30–32^. The difference observed between wt_18_ and wt_25_ is probably reflected by physiological adaptation to colder temperatures. Development or acclimation to 15°C increases the tolerance to colder temperatures by lowering the Na^+^ concentration in the haemolymph, preventing the loss of haemolymph water caused by Na^+^ imbalance at cold temperatures ^33^. Ion imbalance has been proposed as one of the factors that may generate cold-trapping since extracellular and intracellular ion concentrations influence the excitability of muscles and neurons ^34^. The differences see in wt_30_ adults cannot be excluded to be of similar origin, but it has to be noted that these flies were very slow, died quickly, had very few offspring and large quantities of parental lines were needed to acquire enough adults. It is therefore more likely that the results reflect a general deficiency in behaviour. Nonetheless, they give insight into the temperature behaviour of *Drosophila* under extreme hot T_R_.

To test if IGLOO could isolate behavioural phenotypes of canonical mutants we tested a number of fly strains with impaired temperature sensation. Adults carrying the *dtrpA1*^*ins*^ mutation showed a slight increase in mean T_P_ compared to wild-type. This can be explained by the surprising shift in cold avoidance, which increased the range of tolerated temperatures significantly (Fig. 4E, F). This behaviour was not described before. In the original publication by Hamada et al., however, it can be observed that a larger proportion of *dtrpA1* ^*ins*^ than of wild type flies prefer a temperature range of 20°C to 22°C. The authors pooled the data for temperatures from 18°C to 22°C and thereby possibly obscuring differences in avoidance. This explains why in their case avoidance of cold regions near the preferred regions was not different between groups, whereas in the present study, *dtrpA1* ^*ins*^ mutants distribute at slightly cooler temperatures.

In contrast to *dtrpA1*^*ins*^ mutants, HC-ablated flies had a clear T_P_ shift. TRPA1 is reported to be present in internal sensory neurons which are located in the brain ^5,35^, while HCs are external sensors ^8,9,36^. It is easy to imagine that a system that cannot judge its desired temperature has a broader tolerance, contrary to a system with defect sensor that might settle for the wrong temperature. This hypothesis is further corroborated by the larvae experiments. *dtrpA1*^*ins*^ in larvae has been implicated in the sensation of temperatures warmer than the preferred 18°C by body-wall neurons ^11,35,37^. The median T_P_ of *dtrpA1*^*ins*^ larvae is significantly increased, while their tolerance is not (Fig. 4H,I). This resembles the adult HC phenotype (cmp. Fig. 4E,F,H,I).

The difference in T_P_ between late 3rd instar larvae and adults is still present after correction with our model and it is consistent with previous reports ^11,15,38^. In addition, the shift to warmer temperatures in *dtrpA1* ^*ins*^ larvae and the changes in T_P_ obtained when ablating or silencing hot and cold cells in the adult arista ^8^ were also present after the cold bias was removed. These examples demonstrate that our model is a suitable tool to study T_P_ of small poikilothermic animals and to assess the role of different molecules or groups of cells in thermosensation.

Using temperature gradients in addition to two-choice assays has become a standard method, in particular when investigating gene and protein function. IGLOO can help with planning such experiments in several ways: (1) determining temperature extremes for both two-choice assays and gradients, (2) thereby avoiding cold-trapping of the flies, (3) building null-hypotheses for a given temperature experiment, (4) planning the optimal duration of an experiment in order to reach equilibrium, and (5) obtaining avoidance, preferences, and tolerances, not just one preference. By integrating data from adult and larval *Drosophila*, as well as flies that were reared at different temperatures, our model can be applied to a wide range of experimental designs. Furthermore, IGLOO can increase the comparability between studies that use similar methods but differ in essential details such as gradient boundaries or rearing temperatures. Taken together, we anticipate that IGLOO will be a valuable tool for planning experiments and analysing results.

## Methods

### Modified Benzer gravitaxis assay

For the Benzer gravitaxis assay ^39^, a vertical arena with 15 separate lanes was printed using a 3D-printer (Ultimaker 1, Ultimaker BV, Geldermalsen,The Netherlands). The arena was closed to the front with an antiglare acrylic glass pane and inserted into a 3D-printed frame that was open at the top. By sliding the arena up and down in the frame, the flies inside the arena were knocked to the bottom. The Benzer gravitaxis assay was placed inside a temperature-controlled incubator chamber (DigiTherm, Tritech Research Inc., Los Angeles, USA). Temperatures inside the chamber could be set in the range between 6°C and 45°C.

Thirty CantonS wild-type flies of each of the three cohorts (reared at 18° (wt_18_), 25° (wt_25_), 30°C (wt_30_)) were tested per temperature (5 repetitions), and the construction of our Benzer test arena allowed us to test up to 15 flies at the same time. This resulted in over 24 million analysed frames. The flies were sedated by cooling on ice for 30 seconds before being transferred to the Benzer arena, where they were allowed to recover for 10 minutes at room temperature (18°-20°C, thermometer model 383T01, Conrad Electronics, Germany) before testing started. In each trial, the flies were knocked to the bottom of the arena and filmed at 30 frames per second while they walked up to the top, using a Hercules Optical Glass camera (Guillemot Cooperation, La Gacilly, France). Filming was stopped as soon as the last fly reached the top of the arena, or, in temperatures in which flies became less capable of moving, filming was stopped as soon as they had clearly settled at a certain position and did not move further upwards.

### Larva crawling assay

The speed of locomotion of 3^r^^d^ instar CantonS larvae (wt_L_) at different temperatures (8°C-40°C) was tested in the same incubator chamber used for the adults. In total 666 larvae were observed, rendering over 1.4 million frames. A layer of agar (1% agarose) was placed on top of an acrylic glass pane with infrared LED ^40^ inserted into the edges of the pane. Wandering larvae were washed twice in water to remove any food residues and placed on the agar. According to the principle of frustrated total internal reflection ^40^, the infrared light only passes from the acrylic glass to the agar and from the agar to the larva, and is then reflected down to a camera (Teledyne Dalsa Motion Traveller 300, IS – Imaging Solutions GmbH, Eningen, Germany) that recorded the crawling larva from underneath using Troublepix software (NorPix Inc., Montreal, Canada). This resulted in images with increased contrast as most of the light captured by the camera came only from the larva. Once the larva started crawling it was recorded at 50 frames per second for 3 minutes or until it reached the edge of the arena. At very low temperatures where the larvae did not move the recording was started 1 minute after the larva was placed on the agar. Twenty CantonS wild-type larvae were individually tested per temperature (except for 8°C where only 16 larvae were tested). The experiments were carried out in darkness for the larvae as they could not detect the infrared light emitted by the LED used in the setup ^41^.

### Tracing

The traces of adults were extracted using ivTrace (Jens P. Lindemann, Bielefeld University) and analysed in Matlab with respect to walking speed, overall activity, and bout duration. To calculate walking speed of adult flies, we used only upward movements, as falling flies would have resulted in much higher (downward) velocities. Walking speed was then calculated as the difference of the positions in two consecutive frames divided by the frame duration. Sideward movements were ignored. Bout duration was defined as the time span that a fly kept walking upwards without stopping for more than 1 second. The animal was considered active whenever its forward speed was faster than 10% of the body length per second (0.2 mm/s). The activity index is the time spend moving faster than 10% bodylength per second, divided by the time the animal needed to reach the top. We also analysed whether or not a fly crossed the mid height and reached the top of the arena and used this information as a proxy for the overall fitness of the flies.

Larvae were traced with ivTrace (Jens P. Lindemann, Bielefeld University). The crawling speed of wt_L_ was calculated for the entire duration of the recording, and the bout duration was defined as the duration of crawling before stopping for more than 1 second. Activity was determined in the same way as for adults.

### Locomotion in temperature gradient

A temperature gradient was used to determine the T_P_ of larvae and adults. An aluminium slab divided in 5 tracks (dimensions: 50mm × 3mm × 3mm) was used to generate the gradient. The cold side of the gradient was generated by placing a brass cylinder containing ice water with salt on the aluminium slab. The hot side was generated by heating soldering irons. The gradient generated ranged from 14°C (±1°C) to 30°C. For better temperature conductance no agar was used in the arena. Once the gradient was generated a larva or adult was placed into each track, which allowed to obtain individual T_P_ of each animal tested. Adults were anesthetised on ice before being placed in the arena. A translucent acrylic glass covered the tracks to prevent the animals from escaping. The acrylic glass was coated with Sigmacote (Sigma-Aldrich) to prevent the animals from crawling on it. The animals were allowed to walk/crawl and settle in the gradient for 11-12 minutes and were then recorded for 5 minutes at 50 frames per second. The relative humidity was 27% during recordings.

The temperature of the arena was recorded at a rate of 10Hz by 30 thermoresistors (SEMI 833 ET, B+B Thermo-Technik GmbH, Donaueschingen, Germany) that were distributed inside the arena. The signal from the thermoresistors was collected by a program running on an Arduino Micro chip (arduino.cc) and sent via serial communication to a PC running Matlab. This allowed to monitor the gradient online and store the data for further analysis. The traces of the recorded videos were extracted using ivTrace to obtain the position of the animals in the arena over time. This position was then correlated with the temperature recorded for the same position in the arena for every frame, which allowed to obtain the position of each animal in relation to temperature over time.

### Null model for temperature-dependent locomotion - IGLOO

We use a Random Walk Monte Carlo-type (RWMC) model to simulate a fly that has no T_P_ moving in a temperature gradient. IGLOO assumes that flies move randomly within the gradient, but that the speed and bout duration depend on the local temperature. Both parameters were obtained from the Benzer gravitaxis assay and larval crawling data. Further assumptions were that a) the thermal gradient is stable, linear and one-dimensional, b) moving towards the hot or the cold part of the gradient is equally probable at every point in the gradient, and c) the upper and lower borders of the gradient are reflective.

We derived a formula to interpolate the temperature dependent locomotion of *Drosophila* based on the Benzer experiment and larval crawling data (see Equation 1, Fig. 1). This formula consists of three terms: (1) a randomly assigned heading, represented by ±1, (2) a velocity term (in mm/s) *v*(*p, T*_*B*_*T*_*R*_) and (3) a duration term *d(p, T*_*B*_*, T*_*R*_), where *T*_*B*_ is the body temperature, *T*_*R*_ is the rearing temperature and *p* is a probabilistic value. The velocity *v* and the duration *d* are calculated and multiplied, rendering a distance that is added to the position of the animal, resulting in both a new position and a new ambient temperature.

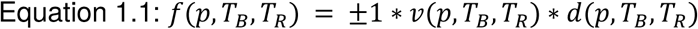

The velocity function can be easily fitted with a two dimensional function consisting of Gaussian distributions (Eq. 2.1). In the temperature domain we need two Gaussians, one with a μ_t1_ of 26 °C and a σ _t1_ of about 5.41 °C and a second one with μ _t2_ = 34 °C and σ_t2_ = 2.27 °C (Eq. 2.2). The probability domain can also be fitted with a Gaussian (μ = −1.49; σ =0.5; Eq. 2.2).

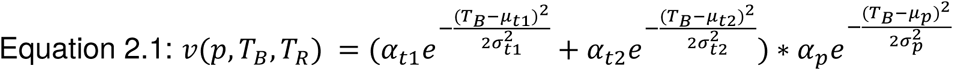

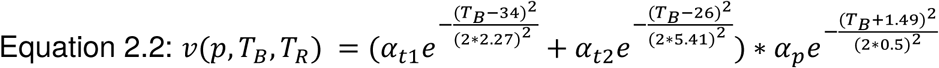

We can fit the influence of the rearing temperature exclusively with the factor α. The fits for α are rather crude using rational or polynomial functions (Eq 2.3) and are only defined within the limits of 18°*C* ≤ *T*_*R*_ ≤ 35°*C*(rearing temperature). As rearing flies below and above these values is unlikely to be successful, this should encompass most encountered rearing temperatures.

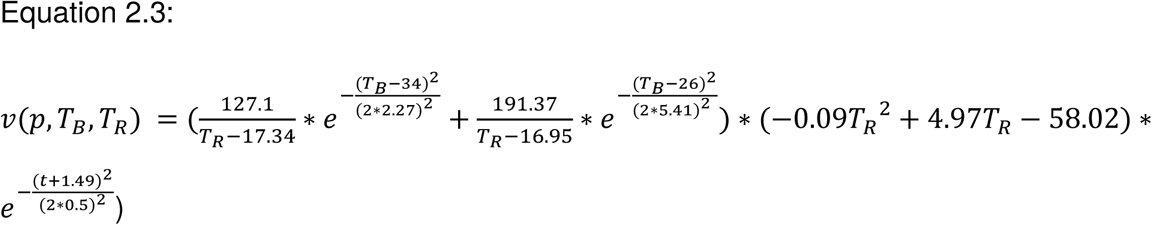

The bout duration function can be denoted as a combination of three Gaussian functions (see equation 2.1) but only needs two rearing dependent factors (*σ*_*t*1_, *μ*_*t*2_). This factors can be easily interpolated with 1^st^ or 2^nd^ degree polynomials:

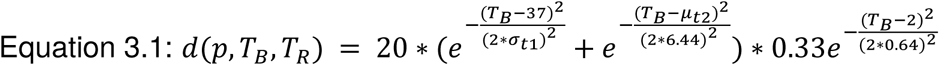

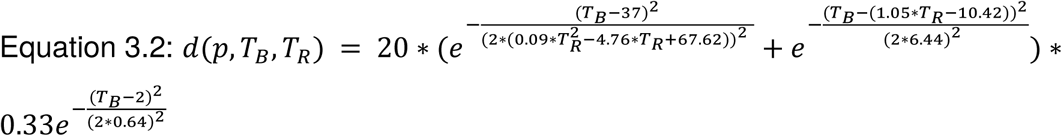

The overall resulting formula is shown in Equation 1.2:

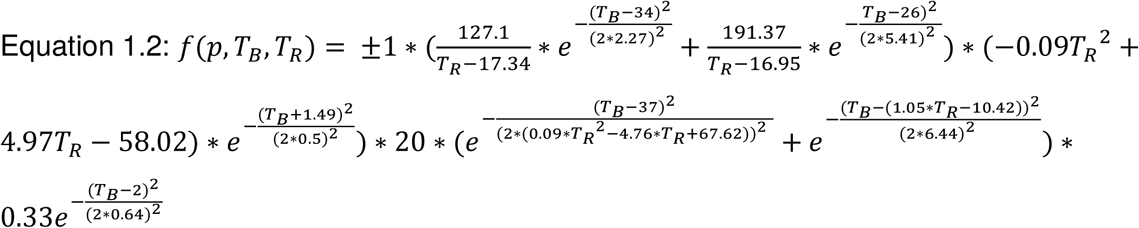

IGLOO simulates larval temperature dependent locomotion with the identical constraints and assumptions as it does adult behaviour, compare Eq 1.1 and 3.1.

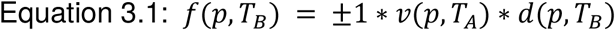

The major adjustments of the algorithm are firstly that is does not include *T*_*R*_, as we only measured data for larvae raised at 25°C. Secondly, larvae exhibited much stronger thermal resilience than adults (see Fig. 2), making adjustments in the interpolation formulae necessary. The larval velocity function is a product of a fourth degree polynomial for the temperature domain and a third degree polynomial for the probabilistic domain (see Eq 3.2). This derives a steady velocity profile that drops off sharply below 10°C.

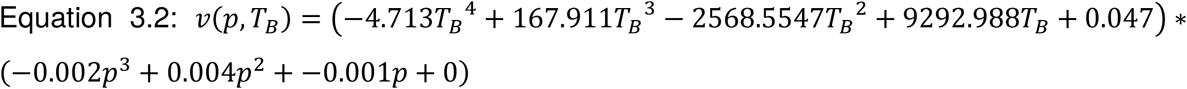

Also the duration function had to be adjusted, as it is now the sum of two products of two Gaussian functions (see Eq 3.3). The values of the temperature dependent Gaussians are 9.0 and 34.0, respectively.

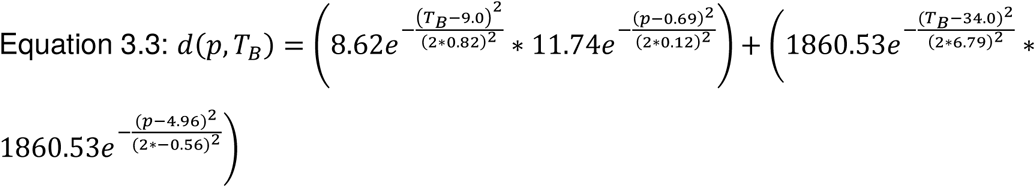

### Thermodynamics of the *Drosophila* body

A simple thermodynamic conduction model, rather than just applying the T_A_ determines the T_B_ of the modelled animal. To estimate the body surface and conductivity roughly, we simulate *Drosophila* as a cylinder of water with a radius of 0.5 mm and a length of 2 mm, resulting in a surface of 7.85 mm^2^. The heat conductance of water is 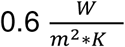 (see ^42^). To determine the change in temperature we used the heat flow formula (see Eq. 4), where *λ* is the heat conductance, *A* is the surface, *D* denotes the wall-to-wall thickness and *T* marks the two different temperatures. In our case, we multiplied the heat flow, which has the unit Joule per second, with the bout duration (*t*) in seconds.

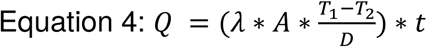

Note that the resulting unit of this formula is now Joule. This energy can be easily factored into a temperature change by taking into account that *Drosophila* weighs about 0.25 mg and 1 Joule heats 1gr of water by 0.2449°K. The assumed cylinder would weigh more (1.57 mg) than a typical *Drosophila*. The cylinder is employed only as rough estimate of the surface available for thermal conduction. This model, however close to the physical process, acts as a low pass filter for T_A_ and could be easily exchanged for one (see Fig. 2B).

### T_P_ correction

We determined the animals’ T_P_ in a gradient by first calculating the probability density of the animal at different temperatures. We tested two different strategies to determine the T_P_ for each individual or a group of flies as a whole. One can either calculate a median temperature histogram from all individual histograms and the 95% confidence interval of each bin or calculate the individual histogram and calculate a confidence of each bin for each individual as rendered by binominal statistics. In both cases we subtract the temperature histogram of the model from the individual or group histogram and check if the 95% confidence interval is above, below or spanning zeros. In the last case this temperature is defined as not preferred, while values above are categorised as preferred and values below as antipreferred. Now one can use a weighted mean to calculate the T_P_, where the temperature bins labels are used to calculate the mean and the bins’ value as weight. Hot and cold antipreference can be calculated, respectively.

### Fly strains

All adult flies and larvae tested in the Benzer gravitaxis and larva crawling assays were of the CantonS wild-type strain and the *brv2*^*MI04916*^, were obtained from Bloomington Stock center (#64349 and #38067, respectively). Larvae were reared at 25°C and adults were reared at three different temperatures: wt_18_ were developed from egg to adult at 18°C, wt_25_ at 25°C, and wt_30_ at 30°C. The *UAS-hid,rpr* line was provided by John Nambu and the *HC-Gal4* and *dtrpaA1*^*ins*^ lines (described by ^8^ and ^5^) were obtained from Paul Garrity, and raised at 25°C. All adults and larvae were reared on the same standard food medium and in the same light-dark cycle (12h light, 12h dark).

### Statistic analysis

Significance test were made as indicated either employing Fisher’s permutation test ^43^ or a paired t-test. All p values were corrected with the Benjamini Hochberg false discovery rate detection ^44^ implemented by David Groppe and colleagues ^45^. All statistical calculations were done using Matlab R2012b (The Mathworks, Natwick, USA).

IGLOO is programmed in python 3 using numpy and scipy packages. The source code is available at https://github.com/zerotonin/igloo and an full online version can be accessed on http://igloo.uni-goettingen.de/.

## Acknowledgements

We thank Paul Garrity (Brandeis University, USA) and John Nambu (University of Massachusetts, USA) for sharing their fly strains with us. We thank Christian Spalthoff (Fraunhofer IEEE, Germany) for assistance with the graphical abstract and Martin Göpfert (Göttingen University, Germany) for financial support and constructive comments. We also thank Ralf Heinrich and Thomas Effertz (Göttingen University) for helpful comments on the manuscript.

## Author Contibutions

AA and BG initiated the study. DG, AA and BG wrote the main manuscript text and prepared figures 1–4. All authors designed the model. HG and BG formalised the model and developed the code. AA recorded adult data basis (Fig. 2), DG recorded larval data basis (Fig. 2). DG recorded experimental proof of concept (Fig. 3). IK developed the web application. All authors reviewed the manuscript.

## Data Availability Statement

The source code of the model is available at https://github.com/zerotonin/igloo. An online application of the model can be accessed at http://igloo.uni-goettingen.de/. Original trajectories and measurement will be shared upon request.

## Abbreviations

IGLOO: **I**GLOO is a **G**radient **LO**comotion m**O**del
wt_18_: CantonS adult flies reared at 18°C temperature
wt_25_: CantonS adult flies reared at 25°C temperature
wt_30_: CantonS adult flies reared at 30°C temperature
wt_L_: CantonS larval flies reared at 25°C temperature
T_P_: temperature preference
T_R_: rearing temperature
T_A_: ambient temperature
T_B_: body temperature
CI: confidence interval

